# Extracellular Vesicles Facilitate Large-Scale, Homogenizing Dynamic Exchange of Proteins and RNA Among Cultured Chinese Hamster Ovary (CHO) and Human Cells

**DOI:** 10.1101/2021.08.20.456971

**Authors:** Jessica Belliveau, Eleftherios T. Papoutsakis

## Abstract

Cells in culture are viewed as unique individuals in a large population communicating through extracellular molecules and, more recently extracellular vesicles (EVs). Our data here paints a different picture: the homogenizing effect of large-scale exchange of cellular material through EVs. We show that Chinese Hamster Ovary (CHO) cells dynamically produce and uptake EVs, to exchange proteins and RNAs at large-scale. To visualize the dynamic production and cellular uptake of EVs, we used correlative confocal microscopy and scanning electron microscopy, as well as flow cytometry to interrogate labeled cells. We employed cells expressing fluorescent proteins (GFP, miRFP703) and tagged cells with protein and RNA dyes. Flow cytometry was used to quantify the exchange of cellular RNA between cells through EVs. This EV-mediated dynamic exchange observed in CHO cultures was also observed in cultures of the human CHRF-288-11 cell line and of primary human hematopoietic stem and progenitor cells. This study demonstrates an underappreciated native cell communication and protein/RNA exchange mechanism mediated by EVs spanning cell type and lines, suggesting the proximity of cells in normal and tumor tissues may also result in prolific cellular exchange. This exchange would be expected to homogenize the cell-population cytoplasm and dynamically regulate cell proliferation and cellular state.

## Introduction

Extracellular vesicles (EVs) are nano- to macro-sized vesicles derived from cytoplasmic or endosomal membranes. EVs are formed by most, if not all, mammalian cell lines and primary cells and constitute a major mechanism of cell-to-cell communication between cells both *in vitro* and *in vivo* (C.-Y. Kao & Papoutsakis, 2018; C. Y. Kao & Papoutsakis, 2019; Raposo & Stoorvogel, 2013). Microparticles (MPs) form from the outward budding of the cytoplasmic membrane and vary in diameter between 100- and 1000-nm. Exosomes, 40- to 100-nm in diameter, are derived from multivesicular endosomes (MVEs) that fuse with the cell membrane, releasing exosomes into the extracellular space (J. Jiang, Kao, & Papoutsakis, 2017; Raposo & Stoorvogel, 2013). EVs are highly enriched in small RNAs compared to the parent cell giving EVs the ability to exert regulatory control on target cells. Native EV cargo also includes proteins, mitochondrial DNA, mRNA, lipids, and organelles/organelle fragments (Boudreau et al., 2014; de Jong et al., 2012; Sansone et al., 2017).

Cells use EVs to communicate stress signals, expel toxic by-product accumulation, and send small RNAs to alter the expression profiles of target cells (de Jong et al., 2012; Han & Rhee, 2018; Njock et al., 2015). There is a large body of literature on the biological effects mediated by EVs *in vitro* and, increasingly, *in vivo* (C. Y. Kao & Papoutsakis, 2019). For example, in *in vitro* studies, uptake of mesenchymal stem cell derived EVs by breast cancer cell lines have been linked to dedifferentiation of breast cancer cells towards cancer stem cells (Sandiford et al., 2021). Megakaryocyte derived microparticles (MkMPs) have been documented as selectively targeting hematopoietic stem and progenitor cells (HSPCs) (J. Jiang et al., 2017; J. L. Jiang, Woulfe, & Papoutsakis, 2014), and mesenchymal stem cell derived EVs are selectively taken up by B, NK, and T cells compared to other lymphocyte cells (Di Trapani et al., 2016). Two, among many now, *in vivo* examples include the impact of human MkMPs in increasing platelet production in a murine model (Escobar, Kao, Das, & Papoutsakis, 2020) and the use of mesenchymal stem cell derived EVs in clinical trials to improve kidney function in patients with chronic kidney disease (Nassar et al., 2016).

Chinese hamster ovary (CHO) cells are the most widely used cell host in the biopharmaceutical industry for production of therapeutic proteins (Han & Rhee, 2018; Keysberg et al., 2021). CHO cells are grown in suspension cultures in large scale bioreactors at high cell densities and CHO cell lines are optimized to extend culture lifespan, increase protein production, and limit the accumulation of toxic by-products (Han & Rhee, 2018). Yet very little is known about the biology and impact of EVs on CHO cells in culture. In a recent study (Keysberg et al., 2021), proteomic and small RNA sequencing of CHO EVs harvested from different phases of growth identified differentially expressed proteins and small RNAs. Additionally, the protein and RNA content of the exosomes were compared to the parent cell, identifying proteins and RNA highly enriched in exosomes, contributing to early understanding of the regulatory effect CHO EVs may have in culture. Additionally, (Seras-Franzoso et al., 2021), EVs from CHO cells overexpressing therapeutic enzymes were recently reported to be effective protein delivery vehicles *in vitro* and *in vivo*, improving enzyme delivery to hard to access organs in traditional enzyme replacement therapies. Fluorescent studies of CHO EVs from these overexpressing cell lines were observed *in vitro* to be taken up by primary mouse aortic endothelial cells (Seras-Franzoso et al., 2021). Current literature provides evidence and support for exploring and detailing the dynamic release and uptake of native EVs of CHO and other cells in culture (Han & Rhee, 2018; Keysberg et al., 2021; Seras-Franzoso et al., 2021). Such detailing would shed light to the potential impact of EV exchange between cells on cell fate both *in vitro* and, by extrapolation, *in vivo*.

In this study we aimed to visualize and quantify the extent of EV exchange in culture of CHO cell lines by utilizing correlative confocal and electron microscopy using both fluorescent protein expressing cell lines, lipophilic protein dyes, and fluorescent RNA stains. Correlative confocal and electron microscopy combines the fluorescent capabilities of confocal microscopy to track and quantify individual EVs with the high-resolution imaging of scanning electron microscopy (SEM) to identify individual EVs at the cell surface prior to EV uptake. In addition to documenting the large-scale exchange of cellular material between CHO cells, extensive levels of EV exchange were also observed in a human megakaryoblastic cell line (CHRF-288-11) and in primary hematopoietic stem and progenitor cells (HSPCs). Based on our findings, we hypothesize that the large-scale exchange of EVs between cells and the associated extensive exchange of proteins, nucleic acids, and lipids, would significantly impact the autonomy of individual cells leading to an ensemble behavior distinct from the behavior of individual cells grown in isolation or in very low culture densities.

## Materials and Methods

### Culture of CHO cells, CHRF cells, and human hematopoietic stem and progenitor (CD34^+^) cells (HSPCs)

A recombinant CHO-K1 (Clone A11 from the Vaccine Research Center at the National Institutes of Health) cell line that expresses the anti-HIV VRC01 antibody was cultured in HyClone ActiPro media (Cytiva) supplemented with 6 mM L-glutamine in 125 mL shake flasks at 120 RPM at 37°C in 5% CO_2_. RFP CHO-K1 (Innoprot) cells were adapted to HyClone SFM4CHO media (Cytiva) and for suspension growth. GFP CHO-K1 cells were a gift from Dr. Kelvin Lee and were cultured in HyClone SFM4CHO media (Cytiva) in 125 mL shake flasks at 120 RPM at 37°C in 5% CO_2_. CHRF-288-11 cells and EmGFP CHRF-288-11 (Fuhrken, Apostolidis, Lindsey, Miller, & Papoutsakis, 2008) were cultured in IMDM supplemented with 10% fetal bovine serum at 37°C in 5% CO_2_. HSPCs (CD34^+^ cells) from mobilized peripheral blood were sourced and cultured as previously described (Kao et al. 2018).

### CHRF cells transiently expressing the fluorescent protein miRFP703

CHRF cells were electroporated following manufacturer protocols for the nucleofector kit V (Lonza) with the pLifeAct-miRFP703 plasmid (Addgene #79993), referred to as miRFP, to express the near-infrared fluorescent protein miRFP703 labeling F-Actin. Briefly, 1×10^6^ CHRF cells were electroporated with 4 μg plasmid DNA using U-023 settings (Amaxa) with 2-mm cuvettes.

### Cell staining using the PKH26 and CFDA-SE, and SYTO RNASelect

Cells and MPs were stained with PKH26 red fluorescent stain (Sigma-Aldrich) following the manufacture protocol to stain proteins in cells and MPs. Isolated cells (180*g*, 4 minutes) and MPs (28,000*g*, 30 minutes) by centrifugation were washed three times in 10% BSA after staining to quench unbound stain. Cells and MPs were resuspended in growth media.

CHO cells stained with 20 mM CFDA-SE dye (Invitrogen,) were incubated at 37°C for 20 minutes. CHO cells stained with CellTracker Deep Red were incubated at 37°C for 30 minutes. Cells were washed three times with PBS before resuspended in growth media.

CHO cells stained with 500 nM SYTO RNASelect green fluorescent cell stain (Molecular Probes) were incubated at 37°C for 20 minutes according to the manufacture protocol. Cells were washed three times with PBS before resuspended in growth media.

### Setup of cocultures

Cocultures of CHO cells, CHRF cells, and CD34+ cells were setup as follows. 2 × 10^6^ CellTracker Deep Red stained CHO cells were cocultured with 2 × 10^6^ CFDA-SE-stained CHO cells. 2 × 10^6^ GFP CHO cells were cocultured with 2 × 10^6^ RFP CHO cells. CHRF cells on day 3 of culture were electroporated with pLifeAct-miRFP703 plasmid as described above. Electroporated cells recovered for 2 hours at 37°C and 5% CO_2_. 2 × 10^6^ miRFP CHRF cells were cocultured with 2 × 10^6^ EmGFP CHRF cells. CD34^+^ cells on day 3 of culture were cocultured at 37°C 5% O_2_ with isolated EmGFP CHRF MPs or PKH26 stained CHRF MPs at a ratio of 100:1. CD34^+^ cells on day 5 of culture followed the same protocol as day 3, except were cultured at 5% CO_2_. All cocultures began in 100 μL for 1 hour before expanding the culture volume to 1 mL. Coculture samples were collected at 24 and/or 48 hours for confocal microscopy as described below.

### Isolation of Extracellular Vesicles

CHO EVs were isolated from culture media with differential ultracentrifugation from day 3 CHO cultures. Cells were removed from the media by centrifugation at 180*g* for 4 minutes. Cellular debris and apoptotic bodies were removed by centrifugation at 2000*g* for 10 minutes. CHO MPs were isolated at 28,000*g* for 30 minutes by ultracentrifugation (Beckman Coulter Optima LE-80K Ultracentrifugation, SW-28 rotor) and concentrated at 28,000*g* for 30 minutes with the Beckman Coulter Optima MAX Ultracentrifuge (TLA-55 rotor). CHO MPs were resuspended in growth media and used immediately or stored at 4°C overnight.

CHO exosomes were isolated from the supernatant after isolating CHO-MPs by filtering (0.22 μm) and centrifuged at 100,000*g* for 90 minutes (SW-28 rotor, Beckman Coulter Optima LE-80K). Then, CHO exosomes were concentrated with the Beckman Coulter Optima MAX Ultracentrifuge (TLA-55 rotor) at 100,000*g* for 90 minutes. CHO exosomes were resuspended in growth media and used immediately or stored at 4°C overnight.

### Zeta potential and Nanoparticle Tracking Analysis (NTA) of EVs

CHO EVs were counted with Nanoparticle Tracking Analysis (NanoSight NS300, Malvern Panalytical) at a 200-fold dilution in filtered PBS. 5 μL of isolated CHO MPs or CHO exosomes were suspended in 1 mL of PBS and the zeta potential was measured on the Litesizer 500 (Anton Paar) at 25°C.

### Flow Cytometric Analysis of Exosomes and Microparticles

Characteristic protein surface markers of exosomes were evaluated on CHO exosomes and microparticles by flow cytometry. Following manufacturer protocols, Dynabeads Protein G Immunoprecipitation Beads (Invitrogen) were coated in exosome targeting capture antibodies, CD63 (SCBT #55051), CD81 (Cell Signaling #10037S), TSG101 (SCBT #7964), and Annexin A1 (Cell Signaling #3299S). Exosomes and MPs were captured on the antibody coated beads overnight with gentle shaking at 4°C. Bead-bound EVs were isolated with a magnetic separator and detected with the same capture antibodies (CD63, CD81, TSG101, Annexin A1) and stained with a fluorescently conjugated secondary antibody for flow cytometry. Control samples for flow cytometry analysis included antibody coated beads to control for the fluorescence of the beads. Additionally, antibody coated beads were treated with the secondary antibody to control for fluorescent antibodies binding to the beads and for the secondary antibody detecting free capture antibody on the bead surface.

### EM of CHO cells and CHO EVs (SEM and TEM)

CHO cells and CHO EVs for SEM were fixed in 4% EM grade glutaraldehyde overnight on poly-L-lysine coated glass coverslips (Fisher Scientific). Samples were processed for SEM imaging by treating the samples to 1% osmium tetroxide for 1 hour, 3 washes with nanopure water, and a series of ethanol dehydration steps (25%, 50%, 75%, 95%, 100%, 100% ethanol). The dehydration process was finished with a CO_2_ critical point drier (Autosamdri 815A, Tousimis) and samples were coated with 5 nm platinum before SEM (Hitachi S4700 and Apreo VolumeScope) imaging.

CHO EVs for TEM imaging were placed on 400 mesh carbon coated copper grids (Electron Microscopy Sciences) and stained with uranyl acetate for negative staining. EV diameters were measured in ImageJ.

Coculture samples in correlative confocal microscopy and SEM microscopy experiments were seeded onto poly-L-lysine coated coverslips or gridded wells (Ibidi) for 10 minutes, fixed with 4% PFA for 10 minutes, and washed three times with PBS. CD34^+^ cell cocultures were stained with PKH26 (before seeding) or Alexa Fluor 647 Phalloidin (after fixation) to visualize the cell membrane. Coculture samples on coverslips were mounted onto glass slides with SlowFade Diamond Antifade Mountant with DAPI (Invitrogen) for imaging. Cells on gridded wells were stored in PBS for imaging.

In correlative confocal microscopy and SEM studies, after CHO cocultures were imaged with confocal microscopy, samples were processed for SEM imaging as described above.

### Image Processing

Post-microscopy analysis was done in Zen Black (Zeiss) for confocal images and Icy (De Chaumont et al., 2012) for correlative processing using the EC-CLEM plug-in (Paul-Gilloteaux et al., 2017). Widefield images with surrounding cells were used as regions of interest (ROIs) to determine the 2D transformation matrix to account for the difference in sample orientation between the confocal microscope and SEM as well as the inversion of images due to the differences in imaging software. The 2D transformation was applied to maximum intensity images of individual cells to align the confocal images to the SEM images. To account for the three-dimensionality of the SEM images relative to the two-dimensionality of the confocal images, ROIs were determined on the perimeter of the maximum intensity projection image to “wrap” the confocal image onto the SEM image.

## Results

### Characterization of CHO EVs

Characterization of CHO EVs harvested from CHO cultures via differential ultracentrifugation included size distribution, zeta potential, surface markers, and surface morphology (Fig. 1A). Measured by nanoparticle tracking analysis (NTA), the mode size of CHO MPs and exosomes is 186 and 107 nm, respectively (Fig. 1B). The zeta potential (approximating the surface charge of the EVs (Midekessa et al., 2020) of the CHO MPs and exosomes was −13.6 and −13.5 mV, respectively (Fig. 1C). Reported exosome values range from −10.3 mV to −55 mV (Han & Rhee, 2018; Kesimer & Gupta, 2015; Midekessa et al., 2020; Sokolova et al., 2011).

**Figure 1:**
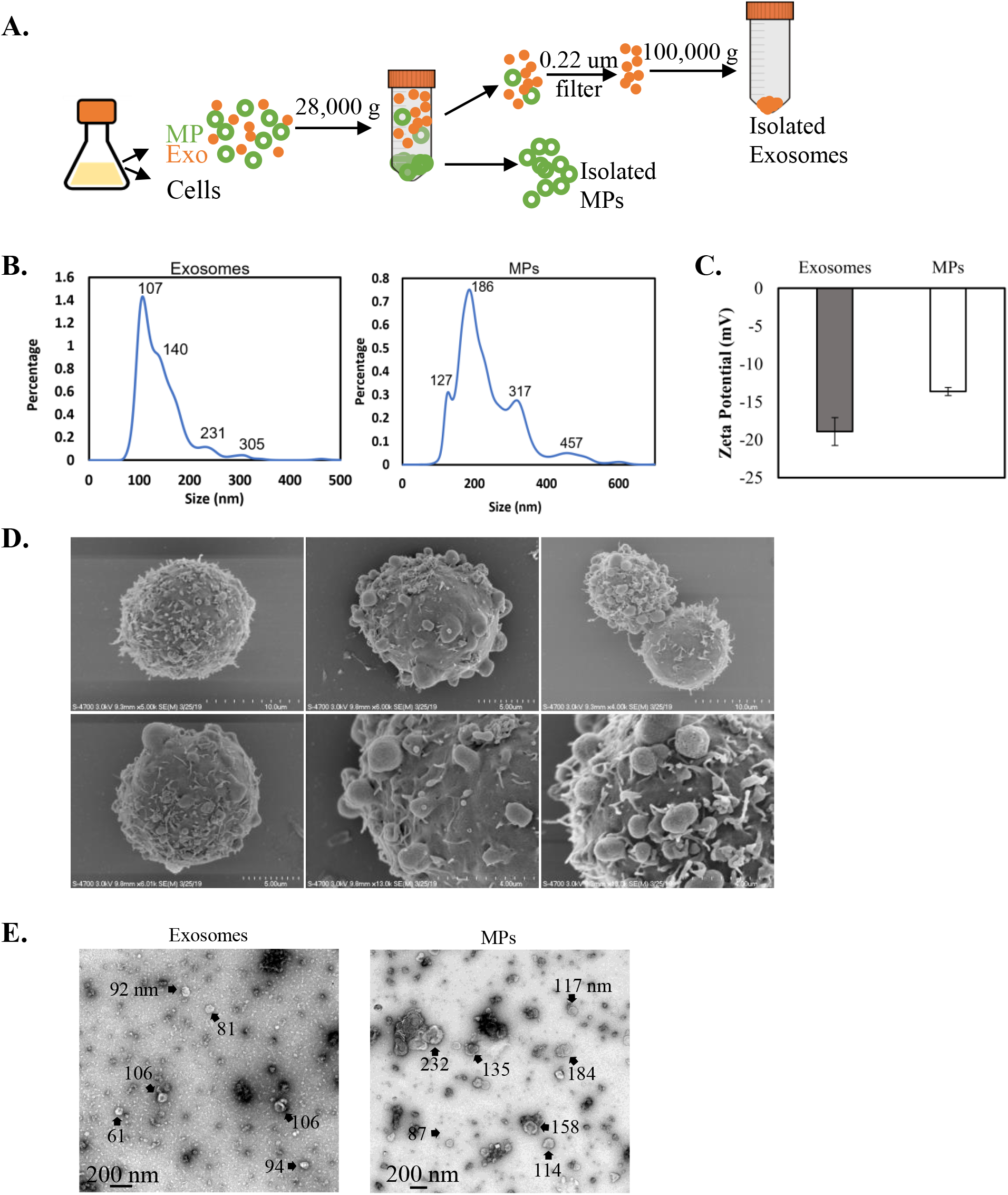
Characterization of CHO EVs. (A) Overview of isolation of CHO exosomes and MPs by differential ultracentrifugation. CHO exosome (left) and MP (right) size characterized by (B) nanoparticle tracking analysis (NTA). (C) Zeta potential of CHO exosomes and MPs. (D) SEM images of CHO cells in culture with EV-like structures at the cell surface. Under normal culture conditions, cells appear with numerous EV-like structures and cells also appear without any EV-like structures at the surface. (E) Size characterization and membrane integrity after ultracentrifugation observed by TEM of CHO exosomes (left) and MPs (right).

Scanning electron microscopy (SEM) and transmission electron microscopy (TEM), was used to examine the surface morphology of CHO EVs (Figs. 1D, E). The diameter of the CHO MPs and CHO exosomes imaged with TEM were measured in ImageJ. The measured EV diameters in the TEM samples were representative of the diameter size distribution measured by NTA. Additionally, EV morphology observed with TEM was similar to other reported CHO EV samples isolated via differential ultracentrifugation (Keysberg et al., 2021). SEM of CHO cells (Fig. 1D) show the surface of CHO cells have small, round, MP-like protrusions that were approximately 0.64- to 1.4-μm in diameter. We hypothesize that some of the protrusions at the cell surface are CHO MPs being produced, and some are CHO EVs the cell is taking up from the culture.

We examined surface protein markers of microparticles, Annexin A1 (Jeppesen et al. 2019), and exosomes, CD63, TSG101, CD81 (Han & Rhee, 2018; Seras-Franzoso et al., 2021), (Fig. 2) to characterize the different isolated EV populations. The mean fluorescent intensity (MFI) of the exosome and MP samples captured on Annexin A1, CD63, TSG101, and CD81 antibody coated beads was compared to the secondary control (n=3). Antibody coated beads were incubated with the secondary antibody without exosomes or MPs in the secondary control. The secondary control determined the baseline for the MFI due to the secondary antibody binding directly to the bead or to the antibodies on the bead surface. Exosomes (Fig. 2A and 2B) had high expression of the surface markers Annexin A1 and CD81, while CD63 and TSG101 were detected on the exosomes at a lower abundance as determined by the differences in MFI. CHO EVs have been reported in other studies to be high in CD81 expression and lower in TSG101 expression (Han & Rhee, 2018; Keysberg et al., 2021). MPs (Fig. 2C) had high expression of the surface markers Annexin A1 and CD81 compared to the secondary control. CD63 and TSG101 were not detected on MPs indicating EV isolation via differential ultracentrifugation isolated two different EV populations. Additionally, Annexin A1 does not appear to be a microparticle specific marker for CHO MPs as it is detected in both the exosome and MP populations.

**Figure 2:**
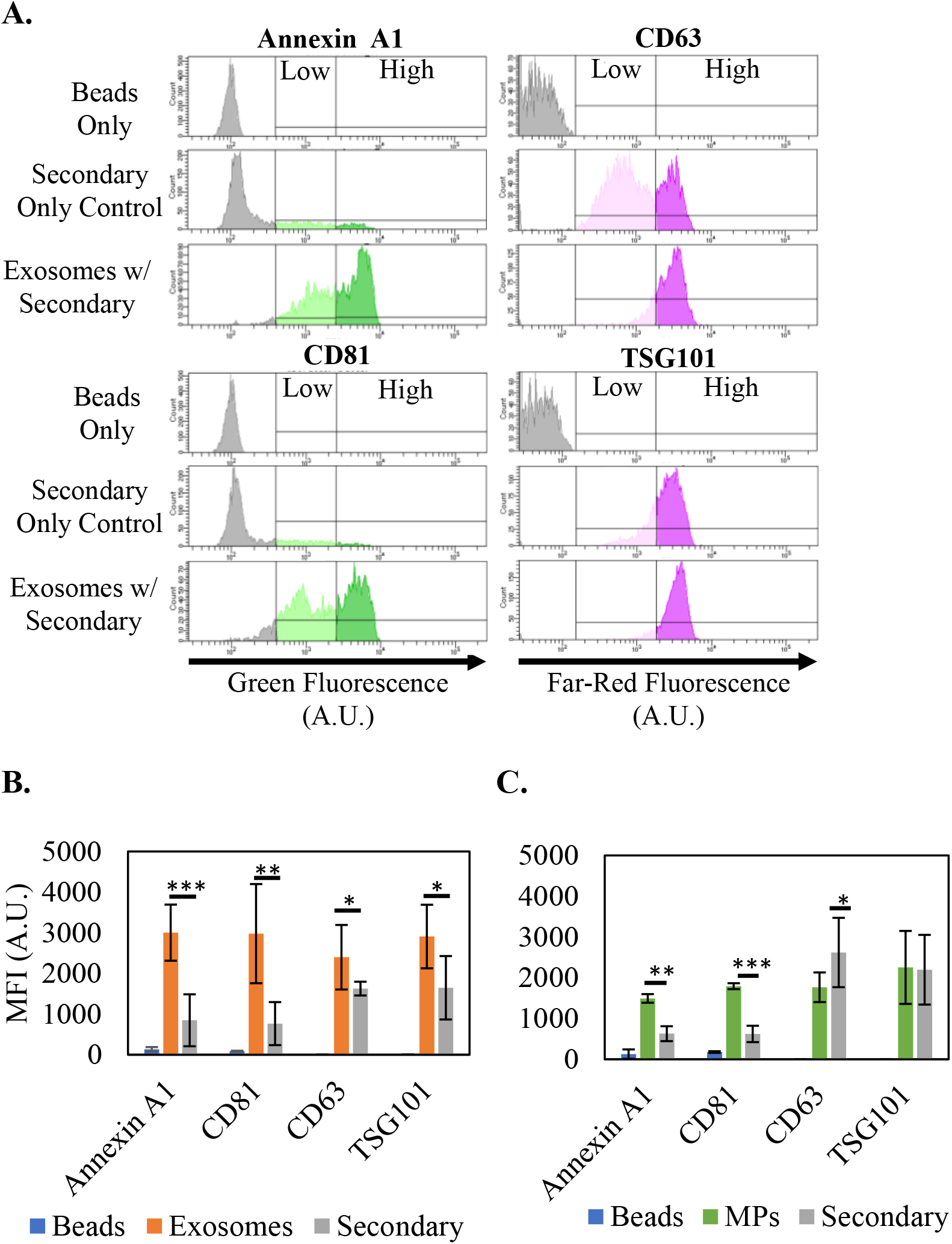
EV surface protein characterization using flow cytometry. CHO exosome and MPs were isolated and captured on antibody-coated capture beads to detect a reported microparticle marker (Annexin A1) and exosome markers (CD63, CD81, TSG101) with flow cytometry. (A) Histograms and (B) mean fluorescent intensity (MFI) of captured CHO exosomes demonstrate the shift in fluorescent intensity of the exosomes with fluorescent secondary antibodies compared to the beads only and the beads treated with the fluorescent secondary antibody. Exosome captured samples display higher levels of fluorescence compared to the low levels in the secondary only control. CHO exosomes were positive for Annexin A1, CD81, and CD63 and had low expression of TSG101. (C) CHO MPs were positive for Annexin A1 and CD81. Error bars represent the standard deviation of three replicates with * p<0.1, ** p<0.05, and ***p<0.01.

### EVs Mediate Profound Dynamic Exchange of Protein and RNA Between CHO Cells

EVs are packaged with RNAs and proteins from the parent cell and are released into the culture. Here we examined the extent to which CHO cells exchange cellular material by releasing and taking up EVs. Correlative confocal and SEM as well as flow cytometry were used to visualize and quantify the extent of material exchange through fluorescently stained total cellular protein and RNA or using cell lines producing fluorescent proteins. Correlative confocal microscopy and SEM combines fluorescent imaging with confocal microscopy and high-resolution imaging via SEM. The combination of these two microscopy methods makes it possible to identify individual EVs (produced by other cells in the culture) on the surface of individual CHO cells as part of the uptake process. Additionally, maximum intensity projection images via confocal microscopy alone enables the visualization of the total number of EVs taken up by a CHO cell while still largely intact in the cell at the time of sampling. This allows for the assessment of the total amount of cellular material being shared amongst cells in culture at a given timepoint.

#### Cell staining using viable protein dyes

CHO cells stained with CellTracker Deep Red, a far-red fluorescent dye, and CFDA-SE, a green-fluorescent dye, were cocultured for 24 hours. The extent of EV exchange between Deep Red stained and CFDA-SE-stained cells was measured with confocal microscopy and correlative confocal/SEM (Fig. 3) and flow cytometry (Suppl. Fig. S1).

**Figure 3:**
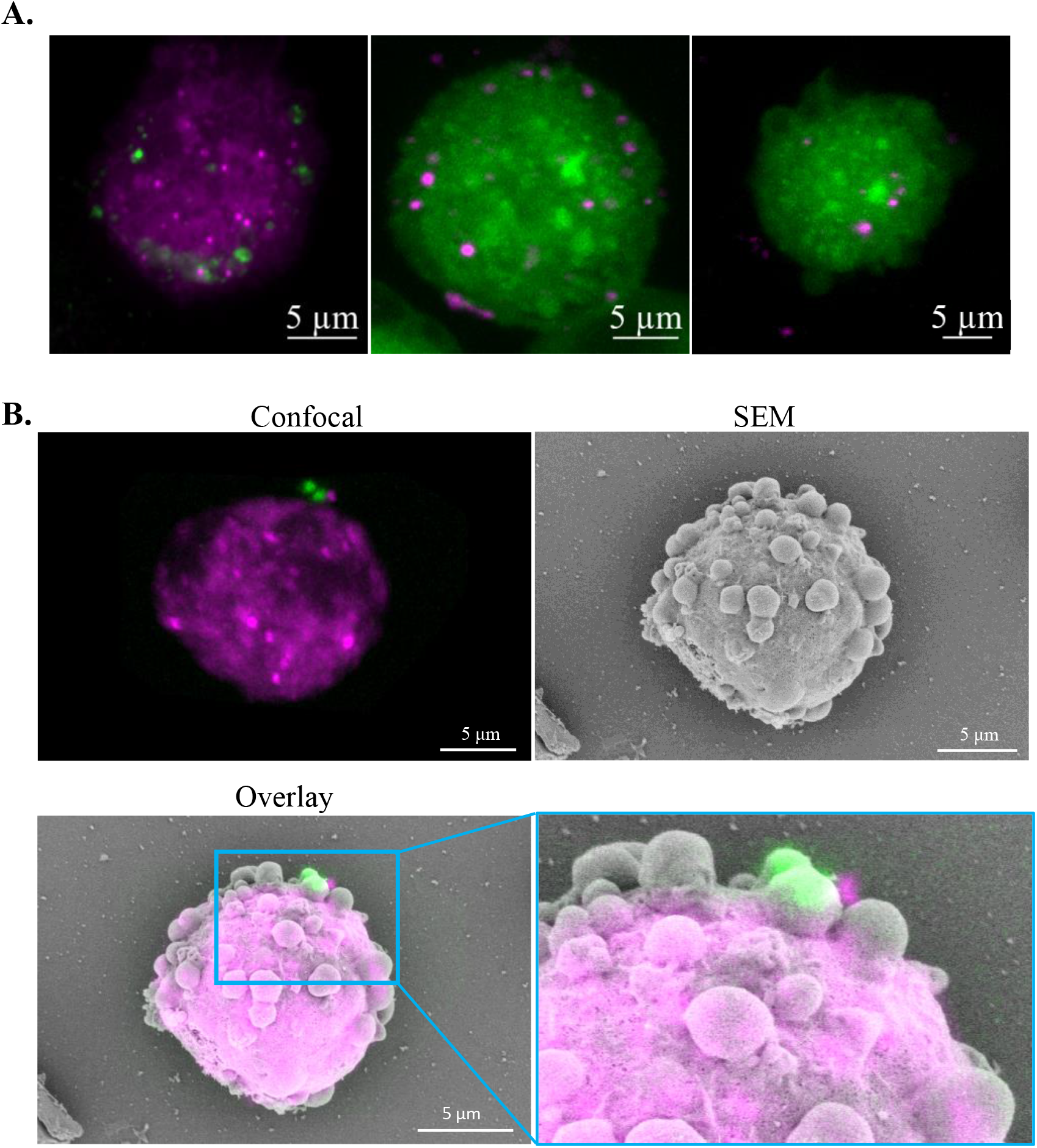
Visualizing the extent of CHO EV exchange between cells using protein stains and correlative microscopy. CFDA-SE stained (green) CHO cells (2 × 10^6^ cells) and CellTracker Deep Red (red) CHO cells (2 × 10^6^ cells) were cocultured in 100 μL growth media for the first hour to enhance the contact between cells and EVs. After the first hour, the coculture was diluted with growth media to 1 mL for 24 hours. At t = 24 hours, cells were assessed for EV exchange. (A) Confocal images. (B) correlative SEM and confocal microscopy identified individual CFDA-SE CHO EV uptake events by a CellTracker DeepRed cell. Two individual CFDA-SE EVs are observed at the cell surface prior to EV uptake.

Maximum intensity projection images, provides a snapshot of representative cells in the coculture at 24 hours is shown in Fig. 3A. In this snapshot, Deep Red stained CHO cells contain EVs from CFDA-SE stained cells and CFDA-SE stained CHO cells contain EVs from Deep Red stained cells. Additionally, the EV-like structures observed at the cell surface in Figs. 1F and 3B are seen in the confocal images with EV-like structures containing stained protein. Correlative confocal/SEM was used to observe specific uptake events on the surface of target CHO cells by identifying fluorescent CHO EVs at the cell surface with confocal microscopy and imaging the cells with SEM to visualize how the CHO EV was interacting with the cell surface (Fig. 3B). In Fig. 3B, a Deep Red stained CHO cell was observed taking up two apparent green stained EVs through membrane fusion.

Flow cytometry (Suppl. Fig. S1) analysis of the coculture began with 53% CFDA-SE positive cells and 53% Deep Red positive cells. At the beginning of culture, approximately 3% of cells were determined to be positive for both fluorescent signals, thus resulting in the 53% positive figure. Gates for CFDA-SE positive cells was determined from pure cultures of unstained cells and Deep Red cells. Gates for Deep Red positive cells were determined from pure cultures of unstained cells and CFDA-SE cells. At 3 hours of coculture, approximately 5% of the population were positive for both stains and this increased to 11% at 24 hours of coculture. Decreases in fluorescent intensity due to cell division and dilution of protein stain was a major challenge in tracking EV exchange by flow cytometry. In stained cells, evaluating the shift in increasing fluorescent intensity as EVs are taken up over culture lifespan competes with the decreasing shift in fluorescence of cells. Additionally, we hypothesize that after an EV is taken up and the cargo is released into the cell, the stained proteins become too dilute to detect by flow cytometry and confocal microscopy. Thus, while fluorescent protein stains are a good tool for understanding EV release and uptake during cell culture, to better understand the extent of EV exchange, CHO cell lines expressing GFP and RFP were utilized. Fluorescent proteins stably expressed in CHO cells do not dilute in fluorescent intensity due to culture growth like cultures using staining with fluorescent dyes.

#### Use of fluorescent-protein expressing CHO cells lines

Cocultures of GFP expressing CHO cells and RFP expressing CHO cells demonstrate the widespread exchange of CHO EVs in culture and quantification of the exchange. CHO EV exchange was quantified with confocal microscopy and flow cytometry (Fig. 4 and Suppl. Fig. 2). Quantification of CHO EV exchange was measured with confocal microscopy by evaluating individual cells in a widefield image (Fig. 5). In two representative widefield images, images of individual CHO cells showed every cell had taken up fluorescent CHO EVs, indicating that CHO EV exchange in CHO cultures is widespread. Confocal microscopy images (Figs. 4A and 5) also demonstrated the range of EV uptake by CHO cells, with some cells containing one or two EVs detectable at a given timepoint and others containing upwards of fifteen EVs. As EVs are taken up, the content of the EVs are released into the cells leading to a decrease in EV-fluorescence intensity and reduced detection. Correlative microscopy revealed individual EV uptake events at the cell surface (Fig. 4B and C). Fig. 4B shows the confocal and SEM images of a GFP CHO cell with RFP CHO EVs and the overlay of the confocal images with the SEM image. Correlating the confocal images with the SEM image resulted in identifying three RFP CHO EVs at the cell surface. SEM images show the GFP CHO cell microvilli interacting with the RFP CHO EV (Fig. 4C). Interactions between EVs and microvilli in the uptake process was also observed by Jiang et al. (Jiang et al. 2017) and suggested that early interactions between EVs and microvilli are important for uptake.

**Figure 4:**
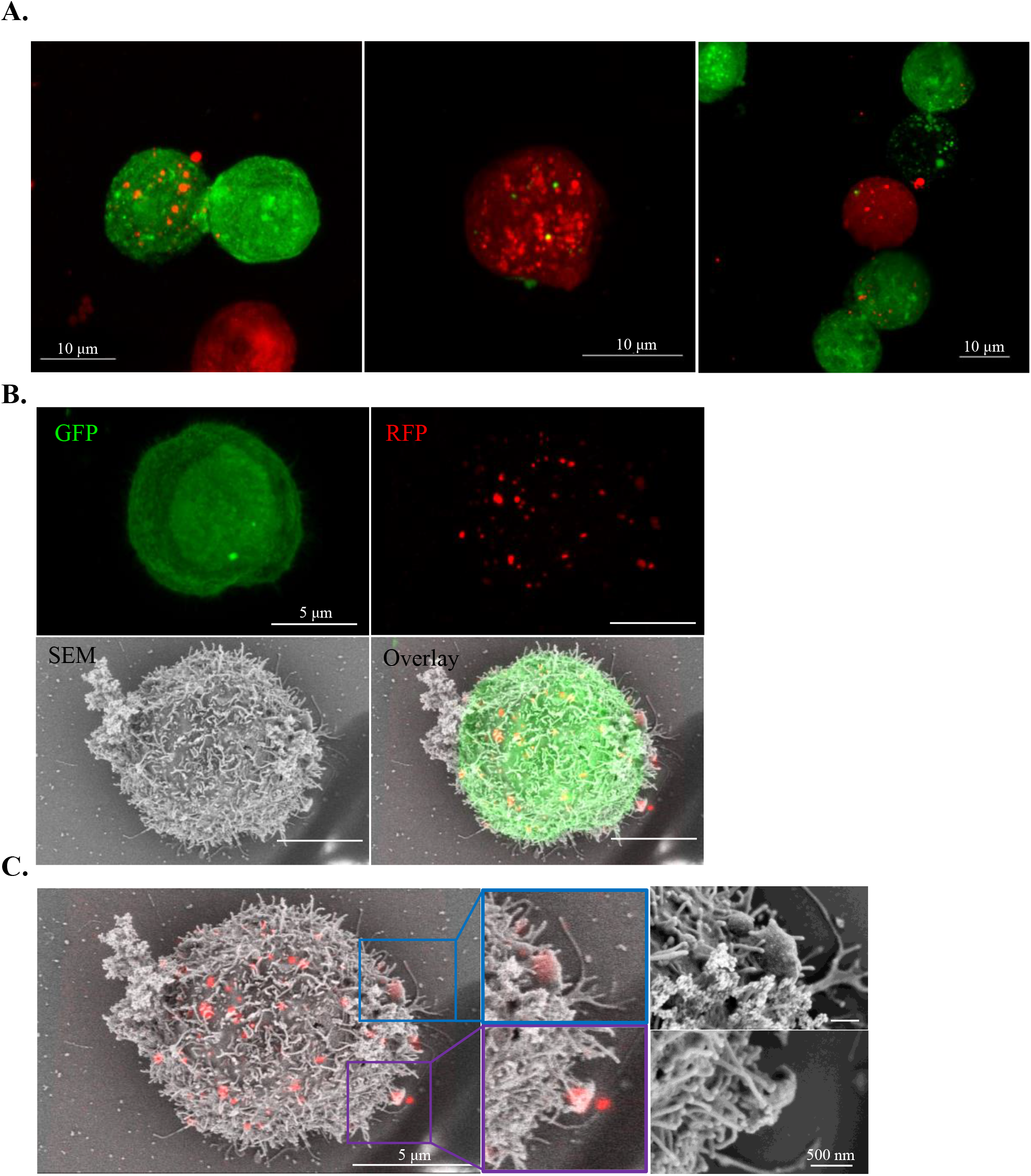
Visualizing the extent of GFP/RFP CHO EVs exchange between cells using correlative microscopy. GFP (green) CHO (2 × 10^6^ cells) and RFP (red) CHO (2 × 10^6^ cells) cells were cocultured in 100 μLgrowth media for the first hour to enhance the contact between cells and EVs. After the first hour, the coculture was diluted with growth media to 1 mL for 24 hours. At t = 24 hours, cells were assessed for EV exchange. (A) Maximum intensity projection confocal microscopy images demonstrate the exchange of EVs in a cell through visualization of intact EVs taken up by cells. (B) Correlative confocal microscopy (GFP and RFP separated into individual channels) and SEM of a GFP cell with RFP CHO EVs at the cell surface. (C) Merged SEM and confocal microscopy (RFP channel only) identify several RFP EVs at the cell surface and high-resolution SEM images of the identified EVs. High-resolution SEM images reveal EVs are interacting with microvilli on the cell membrane potentially as an early cellular mechanism to initiate EV uptake.

**Figure 5:**
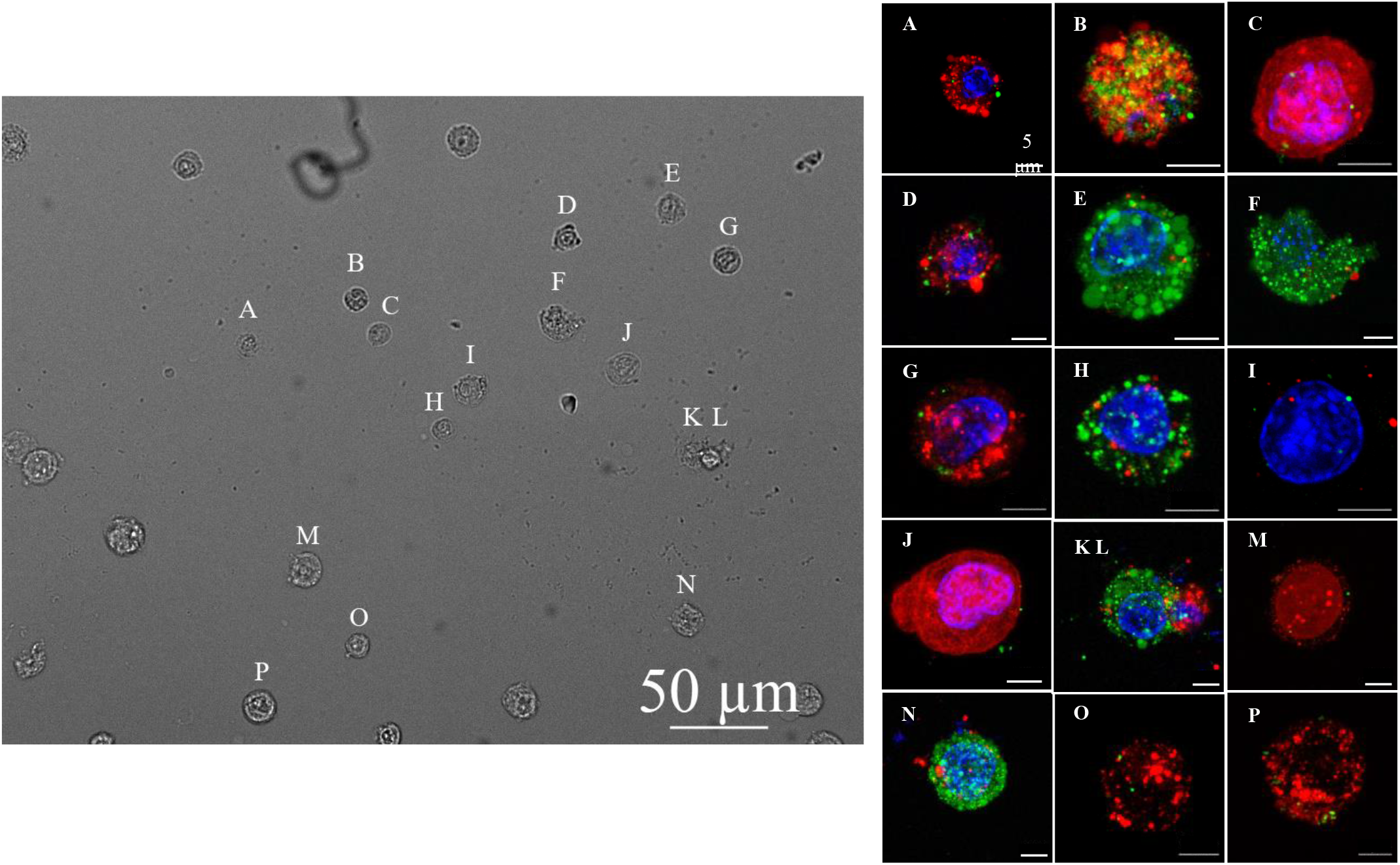
Widefield examination to detail uptake of EVs by CHO cells. Widefield study for CHO EV exchange quantification in a coculture of GFP (green) CHO and RFP (red) CHO cells at 24 hours (DAPI, blue). All cells in the widefield image were positive for GFP and RFP demonstrating the widespread exchange of CHO EVs in culture.

### Staining Cellular RNA via the SYTO RNASelect Dye Enables Flow-Cytometric Demonstration of the EV-Mediated Massive Exchange of Cellular Material Between Cells

To overcome the limitation of using protein stains or fluorescent proteins for flow-cytometric analysis of cellular-material exchange between cells through EVs, we used a specific fluorescent RNA stain. The SYTO RNASelect dye is specific for RNA detection and only fluoresces green when bound to an RNA molecule. Cocultures were designed with half the CHO cells stained with the SYTO RNASelect dye and half the CHO cells unstained. In quantifying CHO EV exchange with flow cytometry (Fig. 6A), 75% of CHO cells were positive for SYTO RNASelect stain after 6 hours and 98% after 24 hours. Confocal microscopy of the cocultures identified individual CHO EVs within targeted CHO cells at 24 hours (Fig. 6B). Similar to the fluorescent protein studies, confocal microscopy can only capture intact EVs taken up by a cell at a given timepoint and does not capture the total number of EVs taken up by a cell throughout the lifespan of the culture. The exchange of EVs observed in the coculture of unstained and SYTO RNASelect stained cells mirrored the levels of exchange observed with confocal microscopy and fluorescent proteins. We hypothesize EV exchange in coculture was observed with the SYTO RNASelect stained cells due to the large abundance of RNA (notably rRNA) in the EVs and the brighter fluorescence intensity of the stain. Flow cytometry analysis of cocultures with fluorescent proteins (Suppl. Fig. 1 and 2) did not capture the massive exchange observed with confocal microscopy, thus suggesting that flow cytometry is not sensitive enough to detect EVs with fluorescent proteins taken up by cells.

**Figure 6:**
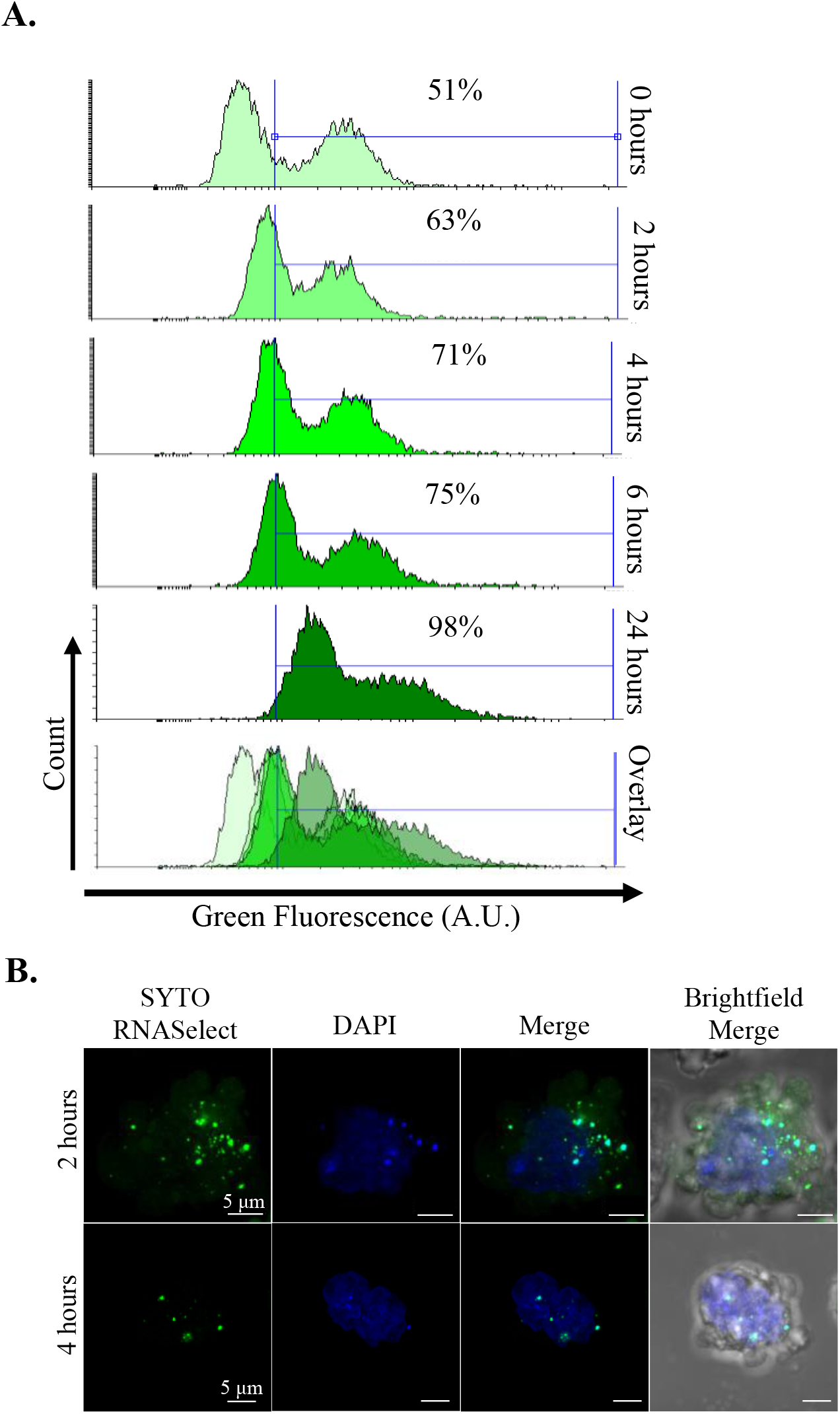
Exchange of cellular RNA between CHO cells through dynamic EV formation and uptake. CHO coculture with half unstained cells and half SYTO RNASelect stained cells. (A) Flow cytometry of the coculture over 24 hours. (B) confocal microscopy of cells containing SYTO RNASelect stained CHO EVs from initially unstained cells after 2 and 4 hours of coculture.

### Dynamic Exchange of Protein Material in the Human CHRF cell line and Primary Human Hematopoietic Stem Cells

Widespread exchange of EVs in culture was also observed in the human CHRF-288-11 (CHRF) megakaryoblastic cell line and in human mobilized peripheral blood CD34^+^ cells (Hematopoietic Stem & Progenitor Cells, HSPCs). EmGFP CHRF cells were cocultured with miRFP CHRF cells for 48 hours. Similar to the CHO cocultures, EmGFP CHRF cells produced EmGFP CHRF EVs and miRFP CHRF cells produced miRFP CHRF EVs and the exchange of fluorescent EVs was observed. Representative images of CHRF cocultures demonstrate the widespread exchange of EVs in cultures (Fig. 7) and numerous taken up EVs were taken up by the target cells. We estimate the extent of EVs exchanged observed with confocal microscopy in CHRF cocultures was approximately 80% compared to that observed in CHO cocultures.

**Figure 7:**
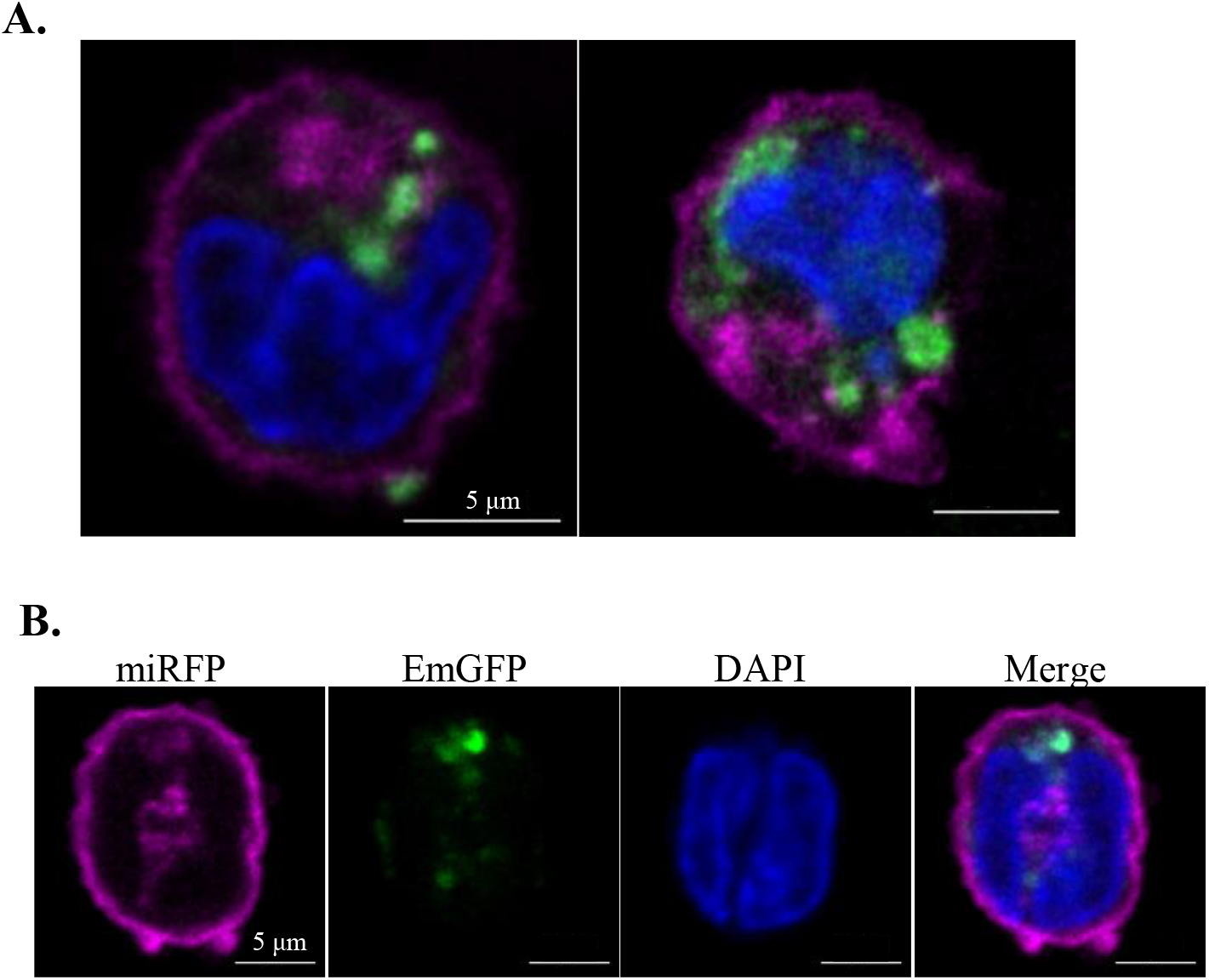
Visualization of EV exchange between CHRF cells in culture. Confocal images of individual cells at the 24-hour time point of a coculture of EmGFP CHRF (green) and miRFP CHRF (magenta) cells. DAPI (blue) is for staining the nuclei. EmGFP CHRF derived EVs are taken up by miRFP CHRF cells. CHRF EVs are observed at the periphery and center of the CHRF cell. (A) In maximum intensity projection confocal images, distinct EmGFP CHRF EV signal is detected in the cytoplasm of miRFP CHRF cells. Fluorescent signal from EVs were both concentrated and diffuse suggesting EVs within the cell are both intact and unloading cargo into the cell. (B) A maximum intensity projection confocal image of a miRFP CHRF cell with both individual fluorescent channels and a merged fluorescent channel. The green channel representing EmGFP CHRF EVs shows intact EVs colocalizing to the cell cytoplasm.

Primary human HSPCs were also examined aiming to demonstrate the generality of cellular material exchange between cells in culture via EVs. It was previously shown that CHRF EVs are taken up by cultured HSPCs (Kao et al. 2018). Human HSPCs from days 3 or 5 of standard HSPC cultures (Kao et al. 2018) were cocultured with either isolated EmGFP CHRF MPs or isolated CHRF MPs stained with the lipophilic membrane stain PKH26 at a ratio of 100 MPs per cell. The HSPC cell membrane was visualized with the green fluorescent actin stain phalloidin and the nucleus was stained with DAPI. Coculture samples of the HSPCs and isolated CHRF EVs were collected at 24 and 48 hours and analyzed with confocal microscopy (Fig. 8). CHRF MPs were readily taken up by day 3 and day 5 HSPCs in coculture. A representative maximum intensity projection image of an HSPC from the coculture (Fig. 8A) demonstrates the total number of intact CHRF MPs at the cell surface and taken up by the cell. HSPC were motile in culture and HSPCs with uropod structures were observed (Fig. 8B). The uropod structure has been reported to be an important mediator and preferred interaction site of EV uptake in HSPCs (J. Jiang et al., 2017).

**Figure 8:**
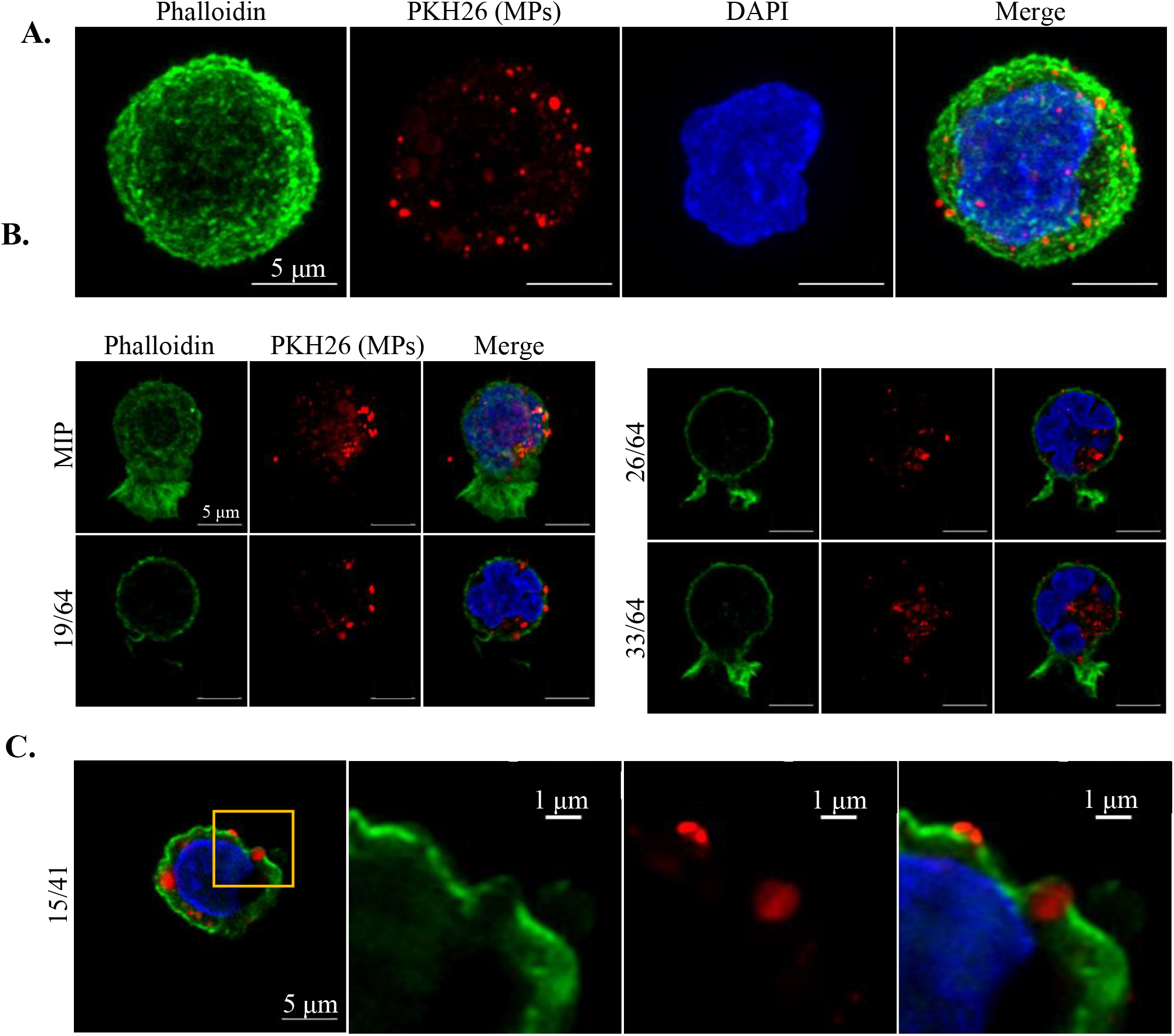
Uptake of CHRF MPs by human HSPCs. Day 3 HSPCs (actin stained with phalloidin, green) were cocultured with CHRF MPs stained with the lipophilic membrane stain PKH26 (red). Confocal images of individual cells at (A) 24 hours and (B,C) 48 hours (nucleus stained with DAPI, blue). (A) Maximum intensity projection (MIP) image of a single HSPC and PKH26 stained CHRF EVs inside the cell. (B) MIP image of a single HPSC containing stained CHRF EVs at the different z-planes of the HSPC. CHRF EVs are observed at the periphery and center of the cell. (C) CHRF EVs are internalized by an HSPC and zoomed-in images of a CHRF MP shortly after uptake at the HSPC membrane.

Early CHRF EV uptake by HSPCs demonstrate the imbedding of the MPs in the cell membrane (Fig. 8C). The cell membrane (actin stain phalloidin, green) wraps around the intact PKH26 stained CHRF EV provides further visual evidence for HSPCs endocytosis of EVs (J. Jiang et al., 2017). Cocultures with CHO, CHRF, and HSPCs demonstrate the widespread phenomena of EV exchange and uptake and represent a mechanism for large amounts of protein and RNA exchange between cells in culture that appears to be independent of cell type.

## Discussion

The cell surface morphology of CHO cells observed via SEM, suggests CHO cells are in a constant process of producing and taking up CHO EVs from the culture. We hypothesize this dynamic EV exchange process allows a culture of suspension cells to share large quantities of protein and genetic material to regulate cellular expression as a whole culture instead of as individual cells.

The exchange of EVs in a culture can be studied with lipophilic protein stains and with fluorescently expressing cell lines. Fluorescent expressing cells lines address concerns of decaying protein stains as cells in culture divide (Wang, Duan, Liu, Fang, & Tan, 2005). Additionally, fluorescent protein expressing cell lines offer improved visualization of EVs on confocal microscopy over lipophilic protein stains. In the situation where fluorescently expressing cell lines are not available, such as primary cell cultures, lipophilic protein stains were shown to be a sufficient tool for visualizing and quantifying the extent of EV exchange in culture. While either approach is suitable when using fluorescent microscopy, flow-cytometric analysis remains a problem with either approach for lack of sensitivity of the fluorescent material associated with the process of EV uptake and dissolution into cells. Indeed, our findings demonstrate vast differences in quantifying EV uptake between flow cytometry and confocal microscopy with the later having greater sensitivity for EV detection. Due to this profound underestimation of EV exchange in culture with flow cytometry, confocal microscopy was the preferred method for observing and quantifying EV uptake and notably the associated exchange of proteins.

Fluorescent RNA specific stains are an alternative method for tracking the uptake of EVs in culture (Li et al., 2014; O’Brien, Breyne, Ughetto, Laurent, & Breakefield, 2020). CHO cells stained with an RNA-specific stain produced EVs containing fluorescent RNA content that was observed transferring between cells. RNA specific stains were easier to detect with flow cytometry compared to protein stains and fluorescent proteins. In a coculture with half the cells unstained and half the cells RNA stained, after 6 hours of coculture, 75% of the cells contained the fluorescent RNA stain via CHO EVs. At 24 hours, 98% of cells in the coculture were positive for the RNA stain. Individual EVs taken up by unstained cells in coculture were observed with confocal microscopy to confirm the unstained cells contained fluorescent CHO EVs. The tracking of EVs via RNA provides quantitative evidence for the vast extent of EV exchange in culture and the kinetics of EV production and uptake in culture. To summarize, in CHO cultures the continuous production and uptake of EVs in culture results in the delivery of proteins and RNA (including regulatory small RNAs but also mRNAs) to any cell from other cells throughout the culture duration. We are beginning to understand the specific protein and RNA content that is exchanged by EVs in CHO cultures and the dynamic regulatory effects the EV cargo has (Keysberg et al., 2021). The impact is this large-scale exchange results in an extensive cytoplasm homogenization of the cell population in culture.

Prior literature has investigated the relationship between cancer cells and stem cells and the transfer of regulatory RNAs via EVs to progress or inhibit tumor growth (Adamo, Dal Collo, Bazzoni, & Krampera, 2019; Fonsato et al., 2012; Lee et al., 2013; Sandiford et al., 2021; Zhou et al., 2019). The ability of cancerous cells and stem cells to communicate throughout the body and in culture presents interesting questions on the effects of how cancer cell derived EVs and stem cell EVs can influence cell expression regulation. These findings combined with the widespread exchange in CHO cells suggests widespread EV exchange is a universal occurrence in eukaryotic cells and cells rely, at least in part, on EVs from other cells to regulate gene expression.

Extracellular vesicles have the potential to spread critical mRNA or regulatory RNAs such as microRNAs, to the cell culture at large. One potential consideration with the extensive exchange of EVs observed in this study is if a spontaneous mutation arises that effects cell viability and proliferation in a way that is detrimental to the production of a recombinant protein or induces apoptosis. The microRNA cluster miR-297-669 has been hypothesized to stimulate apoptosis with nutrient depletion (Aliaksandr Druz et al., 2011) and the miR-466h from this cluster has been studied as it targets anti-apoptotic genes (A. Druz, Betenbaugh, & Shiloach, 2012; Aliaksandr Druz et al., 2011). If a clone arises in a culture that enters apoptosis early and is highly enriched in miR-466h, EVs produced by this cell likely will be enriched in miR-466h. As EVs enriched in miR-466h are circulated in culture and taken up, miR-466h may suppress anti-apoptotic genes and cause the mutated clone to promote apoptosis in the whole culture.

In physiologically relevant studies, extracellular vesicles derived from cells with high-metastatic cancer cells have been observed to promote metastasis in cells with low-metastatic potential both *in vitro* and *in vivo* (Yang et al., 2020). As observed in this study, EVs from the acute megakaryoblastic leukemia cell line CHRF-288-11 were readily taken up by HSPCs in coculture, suggesting cancer cells can influence the expression of stem cells. Studies have reported on the effects of cancer cell derived EVs on stem cells (Chin & Wang, 2016; Daßler-Plenker, Küttner, & Egeblad, 2020; Henrich et al., 2020) and in this study we were able to observe the large extent of cancer-derived EVs uptake by HSPCs.

## Acknowledgements

We thank Sylvain Le Marchand and Deborah Powell and members of the Delaware Biotechnology Institute’s Bio-Imaging Center for assistance with correlative confocal and scanning electron microscopy. We thank Professor Kelvin Lee and Nathaniel Hamaker for the GFP expressing CHO cell line. We thank the Advanced Mammalian Biomanufacturing Innovation Center (AMBIC, NSF Grant1624698), MilliporeSigma, and Sartorius for funding.

## Authorship Contributions

E.T.P. and J.B. designed the study and analyzed the data; J.B. carried out the experiments. E.T.P. and J.B. wrote the manuscript.

## Data Availability Statement

The data used to support the findings of this study are available from the corresponding author upon request.

## References

Adamo, A., Dal Collo, G., Bazzoni, R., & Krampera, M. (2019). Role of mesenchymal stromal cell-derived extracellular vesicles in tumour microenvironment. Biochimica et Biophysica Acta (BBA) - Reviews on Cancer, 1871(1), 192–198. doi:10.1016/j.bbcan.2018.12.001

Boudreau, L. H., Duchez, A.-C., Cloutier, N., Soulet, D., Martin, N., Bollinger, J., … Boilard, E. (2014). Platelets release mitochondria serving as substrate for bactericidal group IIA-secreted phospholipase A2 to promote inflammation. Blood, 124(14), 2173–2183. doi:10.1182/blood-2014-05-573543

Chin, A. R., & Wang, S. E. (2016). Cancer-derived extracellular vesicles: the ‘soil conditioner’ in breast cancer metastasis? Cancer and Metastasis Reviews, 35(4), 669–676. doi:10.1007/s10555-016-9639-8

Daßler-Plenker, J., Küttner, V., & Egeblad, M. (2020). Communication in tiny packages: Exosomes as means of tumor-stroma communication. Biochimica et Biophysica Acta (BBA) - Reviews on Cancer, 1873(2), 188340. doi:10.1016/j.bbcan.2020.188340

De Chaumont, F., Dallongeville, S., Chenouard, N., Hervé, N., Pop, S., Provoost, T., … Olivo-Marin, J.-C. (2012). Icy: an open bioimage informatics platform for extended reproducible research. Nature Methods, 9(7), 690–696. doi:10.1038/nmeth.2075

de Jong, O. G., Verhaar, M. C., Chen, Y., Vader, P., Gremmels, H., Posthuma, G., … van Balkom, B. W. (2012). Cellular stress conditions are reflected in the protein and RNA content of endothelial cell-derived exosomes. J Extracell Vesicles, 1(1), 18396. doi:10.3402/jev.v1i0.18396

Di Trapani, M., Bassi, G., Midolo, M., Gatti, A., Takam Kamga, P., Cassaro, A., … Krampera, M. (2016). Differential and transferable modulatory effects of mesenchymal stromal cell-derived extracellular vesicles on T, B and NK cell functions. Scientific Reports, 6(1), 24120. doi:10.1038/srep24120

Druz, A., Betenbaugh, M., & Shiloach, J. (2012). Glucose depletion activates mmu-miR-466h-5p expression through oxidative stress and inhibition of histone deacetylation. Nucleic Acids Research, 40(15), 7291–7302. doi:10.1093/nar/gks452

Druz, A., Chu, C., Majors, B., Santuary, R., Betenbaugh, M., & Shiloach, J. (2011). A novel microRNA mmu-miR-466h affects apoptosis regulation in mammalian cells. Biotechnology and Bioengineering, 108(7), 1651–1661. doi:10.1002/bit.23092

Escobar, C., Kao, C.-Y., Das, S., & Papoutsakis, E. T. (2020). Human megakaryocytic microparticles induce de novo platelet biogenesis in a wild-type murine model. Blood Advances, 4(5), 804–814. doi:10.1182/bloodadvances.2019000753

Fonsato, V., Collino, F., Herrera, M. B., Cavallari, C., Deregibus, M. C., Cisterna, B., … Camussi, G. (2012). Human Liver Stem Cell-Derived Microvesicles Inhibit Hepatoma Growth in SCID Mice by Delivering Antitumor MicroRNAs. STEM CELLS, 30(9), 1985–1998. doi:10.1002/stem.1161

Fuhrken, P. G., Apostolidis, P. A., Lindsey, S., Miller, W. M., & Papoutsakis, E. T. (2008). Tumor suppressor protein p53 regulates megakaryocytic polyploidization and apoptosis. J Biol Chem, 283(23), 15589–15600. doi:10.1074/jbc.M801923200

Han, S., & Rhee, W. J. (2018). Inhibition of apoptosis using exosomes in Chinese hamster ovary cell culture. Biotechnol Bioeng, 115(5), 1331–1339. doi:10.1002/bit.26549

Henrich, S. E., Mcmahon, K. M., Plebanek, M. P., Calvert, A. E., Feliciano, T. J., Parrish, S., … Thaxton, C. S. (2020). Prostate cancer extracellular vesicles mediate intercellular communication with bone marrow cells and promote metastasis in a cholesterol-dependent manner. Journal of Extracellular Vesicles, 10(2), e12042. doi:10.1002/jev2.12042

Jiang, J., Kao, C. Y., & Papoutsakis, E. T. (2017). How do megakaryocytic microparticles target and deliver cargo to alter the fate of hematopoietic stem cells? J Control Release, 247, 1–18. doi:10.1016/j.jconrel.2016.12.021

Jiang, J. L., Woulfe, D. S., & Papoutsakis, E. T. (2014). Shear enhances thrombopoiesis and formation of microparticles that induce megakaryocytic differentiation of stem cells. Blood, 124(13), 2094–2103. doi:10.1182/blood-2014-01-547927

Kao, C.-Y., & Papoutsakis, E. T. (2018). Engineering human megakaryocytic microparticles for targeted delivery of nucleic acids to hematopoietic stem and progenitor cells. Science Advances, 4(11), eaau6762. doi:10.1126/sciadv.aau6762

Kao, C. Y., & Papoutsakis, E. T. (2019). Extracellular vesicles: exosomes, microparticles, their parts, and their targets to enable their biomanufacturing and clinical applications. Current Opinion in Biotechnology, 60, 89–98. doi:10.1016/j.copbio.2019.01.005

Kesimer, M., & Gupta, R. (2015). Physical characterization and profiling of airway epithelial derived exosomes using light scattering. Methods, 87, 59–63. doi:10.1016/j.ymeth.2015.03.013

Keysberg, C., Hertel, O., Schelletter, L., Busche, T., Sochart, C., Kalinowski, J., … Noll, T. (2021). Exploring the molecular content of CHO exosomes during bioprocessing. Applied Microbiology and Biotechnology, 105(9), 3673–3689. doi:10.1007/s00253-021-11309-8

Lee, J.-K., Park, S.-R., Jung, B.-K., Jeon, Y.-K., Lee, Y.-S., Kim, M.-K., … Kim, C.-W. (2013). Exosomes Derived from Mesenchymal Stem Cells Suppress Angiogenesis by Down-Regulating VEGF Expression in Breast Cancer Cells. PLoS One, 8(12), e84256. doi:10.1371/journal.pone.0084256

Li, M., Zeringer, E., Barta, T., Schageman, J., Cheng, A., & Vlassov, A. V. (2014). Analysis of the RNA content of the exosomes derived from blood serum and urine and its potential as biomarkers. Philosophical Transactions of the Royal Society B: Biological Sciences, 369(1652), 20130502. doi:10.1098/rstb.2013.0502

Midekessa, G., Godakumara, K., Ord, J., Viil, J., Lättekivi, F., Dissanayake, K., … Fazeli, A. (2020). Zeta Potential of Extracellular Vesicles: Toward Understanding the Attributes that Determine Colloidal Stability. ACS Omega, 5(27), 16701–16710. doi:10.1021/acsomega.0c01582

Nassar, W., El-Ansary, M., Sabry, D., Mostafa, M. A., Fayad, T., Kotb, E., … Adel, H. (2016). Umbilical cord mesenchymal stem cells derived extracellular vesicles can safely ameliorate the progression of chronic kidney diseases. Biomaterials Research, 20, 21. doi:10.1186/s40824-016-0068-0

Njock, M. S., Cheng, H. S., Dang, L. T., Nazari-Jahantigh, M., Lau, A. C., Boudreau, E., … Fish, J. E. (2015). Endothelial cells suppress monocyte activation through secretion of extracellular vesicles containing antiinflammatory microRNAs. Blood, 125(20), 3202–3212. doi:10.1182/blood-2014-11-611046

O’Brien, K., Breyne, K., Ughetto, S., Laurent, L. C., & Breakefield, X. O. (2020). RNA delivery by extracellular vesicles in mammalian cells and its applications. Nature Reviews Molecular Cell Biology, 21(10), 585–606. doi:10.1038/s41580-020-0251-y

Paul-Gilloteaux, P., Heiligenstein, X., Belle, M., Domart, M.-C., Larijani, B., Collinson, L., … Salamero, J. (2017). eC-CLEM: flexible multidimensional registration software for correlative microscopies. Nature Methods, 14(2), 102–103. doi:10.1038/nmeth.4170

Raposo, G., & Stoorvogel, W. (2013). Extracellular vesicles: exosomes, microvesicles, and friends. J Cell Biol, 200(4), 373–383. doi:10.1083/jcb.201211138

Sandiford, O. A., Donnelly, R. J., El-Far, M. H., Burgmeyer, L. M., Sinha, G., Pamarthi, S. H., … Rameshwar, P. (2021). Mesenchymal Stem Cell–Secreted Extracellular Vesicles Instruct Stepwise Dedifferentiation of Breast Cancer Cells into Dormancy at the Bone Marrow Perivascular Region. Cancer Research, 81(6), 1567–1582. doi:10.1158/0008-5472.can-20-2434

Sansone, P., Savini, C., Kurelac, I., Chang, Q., Amato, L. B., Strillacci, A., … Bromberg, J. (2017). Packaging and transfer of mitochondrial DNA via exosomes regulate escape from dormancy in hormonal therapy-resistant breast cancer. Proc Natl Acad Sci U S A, 114(43), E9066–E9075. doi:10.1073/pnas.1704862114

Seras-Franzoso, J., Díaz-Riascos, Z. V., Corchero, J. L., González, P., García-Aranda, N., Mandaña, M., … Abasolo, I. (2021). Extracellular vesicles from recombinant cell factories improve the activity and efficacy of enzymes defective in lysosomal storage disorders. Journal of Extracellular Vesicles, 10(5), e12058. doi:10.1002/jev2.12058

Sokolova, V., Ludwig, A. K., Hornung, S., Rotan, O., Horn, P. A., Epple, M., & Giebel, B. (2011). Characterisation of exosomes derived from human cells by nanoparticle tracking analysis and scanning electron microscopy. Colloids Surf B Biointerfaces, 87(1), 146–150. doi:10.1016/j.colsurfb.2011.05.013

Wang, X.-Q., Duan, X.-M., Liu, L.-H., Fang, Y.-Q., & Tan, Y. (2005). Carboxyfluorescein Diacetate Succinimidyl Ester Fluorescent Dye for Cell Labeling. Acta Biochimica et Biophysica Sinica, 37(6), 379–385. doi:10.1111/j.1745-7270.2005.00051.x

Yang, B., Feng, X., Liu, H., Tong, R., Wu, J., Li, C., … Zheng, S. (2020). High-metastatic cancer cells derived exosomal miR92a-3p promotes epithelial-mesenchymal transition and metastasis of low-metastatic cancer cells by regulating PTEN/Akt pathway in hepatocellular carcinoma. Oncogene, 39(42), 6529–6543. doi:10.1038/s41388-020-01450-5

Zhou, X., Li, T., Chen, Y., Zhang, N., Wang, P., Liang, Y., … Chen, H. (2019). Mesenchymal stem cell-derived extracellular vesicles promote the in vitro proliferation and migration of breast cancer cells through the activation of the ERK pathway. International Journal of Oncology, 54, 1843–1852. doi:10.3892/ijo.2019.4747

