## Supplementary Figures for "Extracellular Vesicles Facilitate Large-Scale, Homogenizing Dynamic Exchange of Proteins and RNA Among Cultured Chinese Hamster Ovary (CHO) and Human Cells"

**Supplementary Figure 1: Flow-cytometric examination of CHO EV exchange using protein-staining dyes .** Coculture of CDFA-SE stained CHO cells and CellTracker Deep Red CHO cells to quantify the exchange of individual EVs by flow cytometry at 0, 3, and 24 hours. In each panel, the distributions on the right represent the shifting fluorescent population of the initially stained CFDA-SE (FITC) or CellTracker Deep Red cells (APC). The distributions on the left represent the shift in fluorescent intensity of the initially negative CFDA-SE or CellTracker Deep Red cells due to uptake of fluorescent CHO EVs.

**Supplementary Figure 2: Flow-cytometric assessment of CHO EV exchange when using CHO cells expressing fluorescent proteins.** Flow cytometry time course of a GFP (green) CHO and RFP (red) CHO coculture over 24 hours. Minimal exchange of EVs in the coculture was observed with flow cytometry and was not representative of the levels of exchange observed using confocal microscopy studies.

**Supplementary Figure 1:**

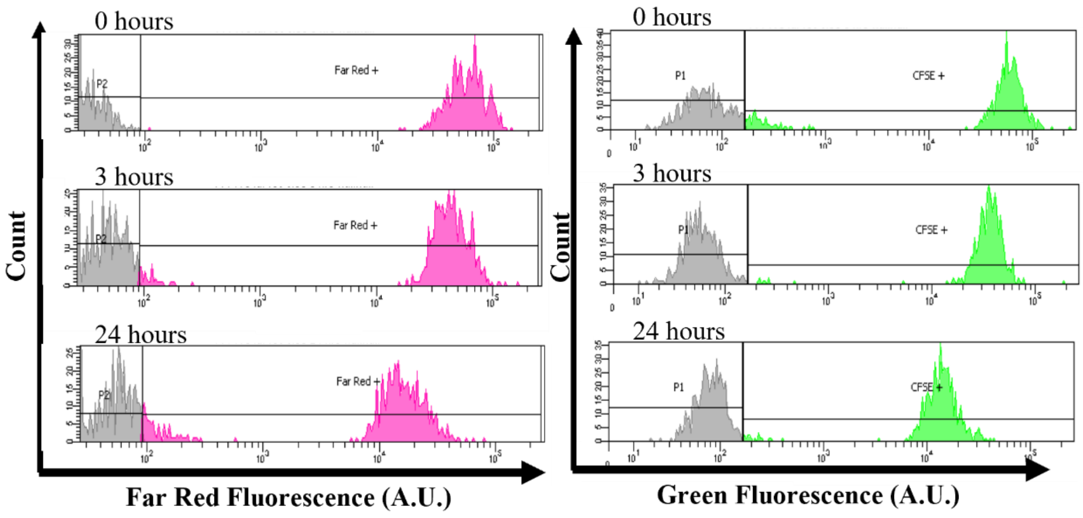


**Supplementary Figure 2:**

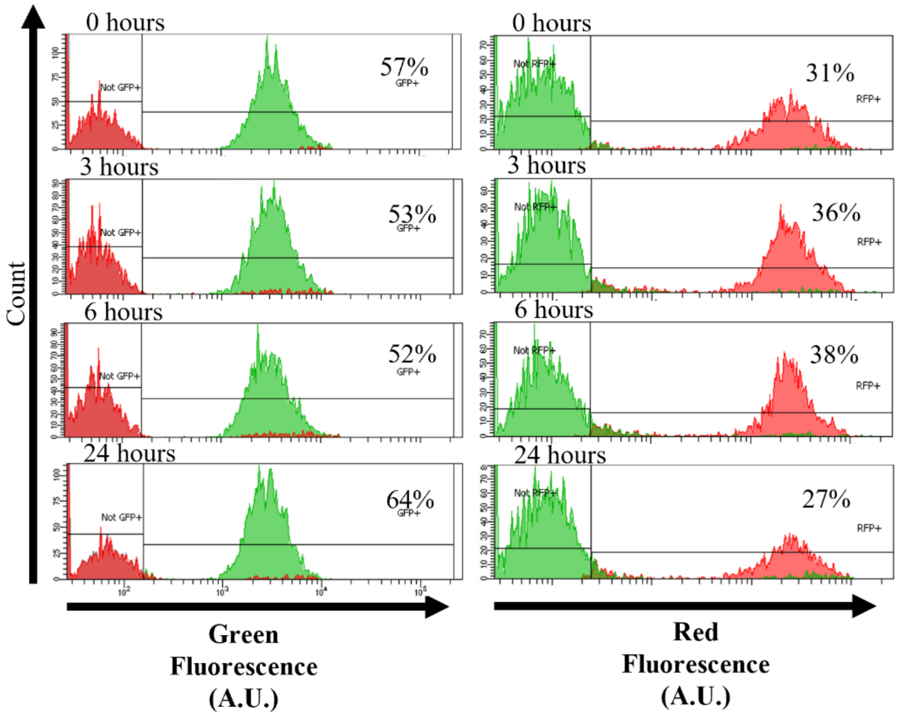
Count
0 hours
57%
GFP+
Not GFP+
3 hours
53%
6 hours
52%
24 hours
64%
31%
RFP+
Not RFP+
36%
38%
27%
Green
Fluorescence
(A.U.)
Red
Fluorescence
(A.U.)
